# The Respiratory Syncytial Virus G Protein Enhances the Immune Responses to the RSV F Protein in an Enveloped Virus-like Particle Vaccine Candidate

**DOI:** 10.1101/2022.09.12.507712

**Authors:** Lori McGinnes Cullen, Bin Luo, Zhiyun Wen, Lan Zang, Eberhard Durr, Trudy G. Morrison

**Affiliations:** Department of Microbiology and Physiological Systems, Program in Immunology and Microbiology, University of Massachusetts Chan Medical School, Worcester, MA; Pharmacology, Merck & Co., Inc., West Point, PA, USA; Infectious Diseases and Vaccines Discovery, Merck &Co., Inc., West Point, PA, USA

## Abstract

Respiratory syncytial virus (RSV) is a serious human respiratory pathogen, but no RSV vaccine has been licensed. Many of the vaccine candidates are focused on the viral F protein. However, it is the G protein that binds the likely receptor, CX3CR1, in human alveolar lung cells raising the question of the importance of the G protein in vaccine candidates. Using virus-like particle (VLP) vaccine candidates, we have directly compared VLPs containing only the pre-fusion F protein, only the G protein, or both glycoproteins. We report that VLPs containing both glycoproteins bind to anti-F protein specific monoclonal antibodies differently than VLPs containing only the pre-fusion F protein. Using RSV naïve cotton rats as an animal model, we have found that VLPs assembled only with the pre-F protein stimulated extremely weak neutralizing antibody (NAb) titers as did VLPs assembled with G protein. However, VLPs assembled with both glycoproteins stimulated quite robust neutralizing antibody titers, titers that were significantly higher than the combined titers induced by pre-F only or G only VLPs. VLPs assembled with both glycoproteins induced improved protection of the animals from RSV challenge compared to pre-F VLPs and induced significantly higher levels of antibodies specific for F protein antigenic sites 0, site III, and AM14 binding site compared with VLPs containing only the pre-F protein. These combined results indicate that assembly of pre-F protein with G protein in VLPs further stabilized the pre-fusion conformation or otherwise altered the conformation of the F protein increasing the induction of protective antibodies.

**Importance:** RSV causes significant disease in infants, young children, and the elderly. Thus, development of an effective vaccine for these populations is a priority. Most ongoing efforts in RSV vaccine development have focused on the viral fusion (F) protein, however, the importance of inclusion of G in vaccine candidates is unclear. Here, using VLPs assembled with only the F protein or only the G protein or both glycoproteins, we show that VLPs assembled with both glycoproteins are a far superior vaccine, in a cotton rat model, than VLPs containing only F protein or only G protein. The results show that the presence of G protein in the VLPs influences the conformation of the F protein and the immune responses to F protein resulting in significantly higher neutralizing antibody titers and better protection from RSV challenge. These results suggest that inclusion of G protein in a vaccine candidate may improve its effectiveness.

## Introduction

Respiratory syncytial virus (RSV) causes acute lower respiratory infections, which are a particularly serious risk for infants, young children, and the elderly. The burden of disease in children is estimated to be 33 million cases/year world-wide with 3.2 million hospitalizations and nearly 200,000 deaths (1, 2). RSV infection impacts the elderly at levels often comparable to influenza infections, resulting in 179,000 hospitalizations/year in the US (3-7).

No RSV vaccine has yet been licensed to date although there is clearly a significant need. Indeed, there are intensive efforts to develop an RSV vaccine (8). While there are two predominant surface glycoproteins in virions, the F (fusion) and the G (glycoprotein) proteins, the sequence of the F protein is more conserved across different strains/serotypes of the virus and is thought likely to induce a broader spectrum of protective immunity than the G protein (9). Thus, most of the vaccine candidates in preclinical and clinical development are focused on inducing immune responses to the RSV F protein and many contain only the F protein (8). This focus on F protein is also due to reports of the association of G protein with enhanced respiratory disease upon RSV challenge (reviewed in Anderson, et al (10)). However, the major receptor for virus entry into human lung cells is thought to be CX3CR1, which binds the G protein (11-14). Thus, antibodies specific for the G protein CX3CR1 binding site should be protective, and, indeed, a monoclonal antibody specific to the G protein is protective in animal models (15). Assays for RSV neutralizing antibodies (NAb) induced by vaccine candidates usually utilize tissue culture cells to titer these antibodies but most commonly used tissue culture cells either do not express CX3CR1 or the virus uses proteoglycans as a receptor (16) which bind both F and G (17, 18). Thus, even in candidates containing the G protein, neutralizing antibody titers directed against the G protein are often not adequately assessed. These considerations raise the question of the importance of anti-G protein antibodies in protection from human disease and, thus, inclusion of the G protein in vaccine candidates.

We have developed virus-like particle (VLP) RSV vaccine candidates based on the Newcastle disease virus (NDV) core proteins M and NP. Our VLPs contain both the RSV pre-F and G protein ectodomains fused to the transmembrane and cytoplasmic domains of the NDV F and HN, respectively, for efficient assembly into the VLPs (19-22). We have shown that these VLPs induce both anti-RSV F and anti-RSV G antibodies and good protective responses in the cotton rat model commonly used in RSV studies (19-23).

Since the majority of humans have experienced multiple RSV infections beginning at two to five years of age (24, 25), we compared responses to VLP immunization of RSV naïve and RSV experienced (RSV primed) animals to mimic the human population. We showed that in both populations the VLPs stimulated much higher neutralizing antibody (NAb) tiers in cotton rats than soluble form of the pre-fusion F protein (22). In RSV naïve animals, VLPs stimulated at least 16- fold higher NAb titers than the soluble F protein, which stimulated barely detectable NAbs. In animals previously infected with RSV (RSV primed), we reported that, while soluble F protein stimulated increased levels of NAbs in this population compared to RSV naïve animals, VLPs still stimulated 8-11 fold higher titers than the soluble protein. The differences in neutralizing antibody titers induced by VLPs vs soluble F protein may be due to the particulate nature of the VLPs (26). However, it also seemed possible that the presence of G protein in the VLPs contributed to the differences in neutralizing antibody titers induced by VLPs and soluble pre-F protein.

Here, we directly addressed the importance of the pre-F and G protein sequences in inducing neutralizing antibodies by comparing antibody responses in RSV naïve cotton rats immunized with VLPs assembled only with the RSV pre-fusion F protein or only the RSV G protein or with both proteins. We report here that VLPs assembled with only the pre-fusion F protein stimulated very weak neutralizing antibody titers. VLPs assembled with G protein stimulated only slightly higher titers. However, VLPs assembled with both the pre-fusion F and G protein sequences stimulated quite robust neutralizing antibody titers, titers than were significantly higher than the combined titers of NAbs induced by pre-F only or G only VLPs. Consistent with these observations, we report that animals immunized with VLPs containing only the pre-F protein were less protected from RSV challenge than animals immunized with VLPs containing both pre-F and G proteins or VLPs that contain only G protein. Our results further suggest that assembly of the F protein in VLPs in the presence of G protein influences the conformation of the F protein.

## Results

### Preparation and Properties of VLPs

To assess the role of RSV proteins F, G, or a combination of both in induction of protective responses in cotton rats by a VLP vaccine candidate, three different VLPs were assembled with the NDV M and NP core proteins. One VLP contained the mutation stabilized pre-fusion F protein, DS Cav1 (27), while the second contained the G protein. The third was assembled with both glycoproteins. The RSV glycoproteins were assembled into VLPs as chimera proteins. The ectodomain sequences of the RSV pre-F protein were fused to the foldon (28) sequence and the NDV F protein transmembrane and cytoplasmic domains to produce the pre-F/F chimera protein (29). The ectodomain of the RSV G protein was fused to the cytoplasmic and transmembrane domains of the NDV HN protein to produce the H/G chimera protein (19). VLPs were assembled in and released from avian cells transfected with the cDNAs encoding the RSV F chimera protein (pre-F/F abbreviated as F/F throughout), or the RSV G protein chimera protein (H/G), or both RSV chimera proteins as well as cDNAs encoding the NDV M and NP proteins. VLPs were purified from cell supernatants, quantified, and validated as previously described (19, 22, 30, 31)

The RSV F and G protein content of purified VLPs were quantified by Western blots using anti-RSV F protein or G protein antibodies (22, 32). The three VLP preparations were then adjusted to contain the same F protein content or the same G protein content (Figure 1). Interestingly, the presence of G protein in the VLP preparations was associated with more efficient cleavage of the F protein suggesting an influence of G protein on the F protein conformation.

**Figure 1:**
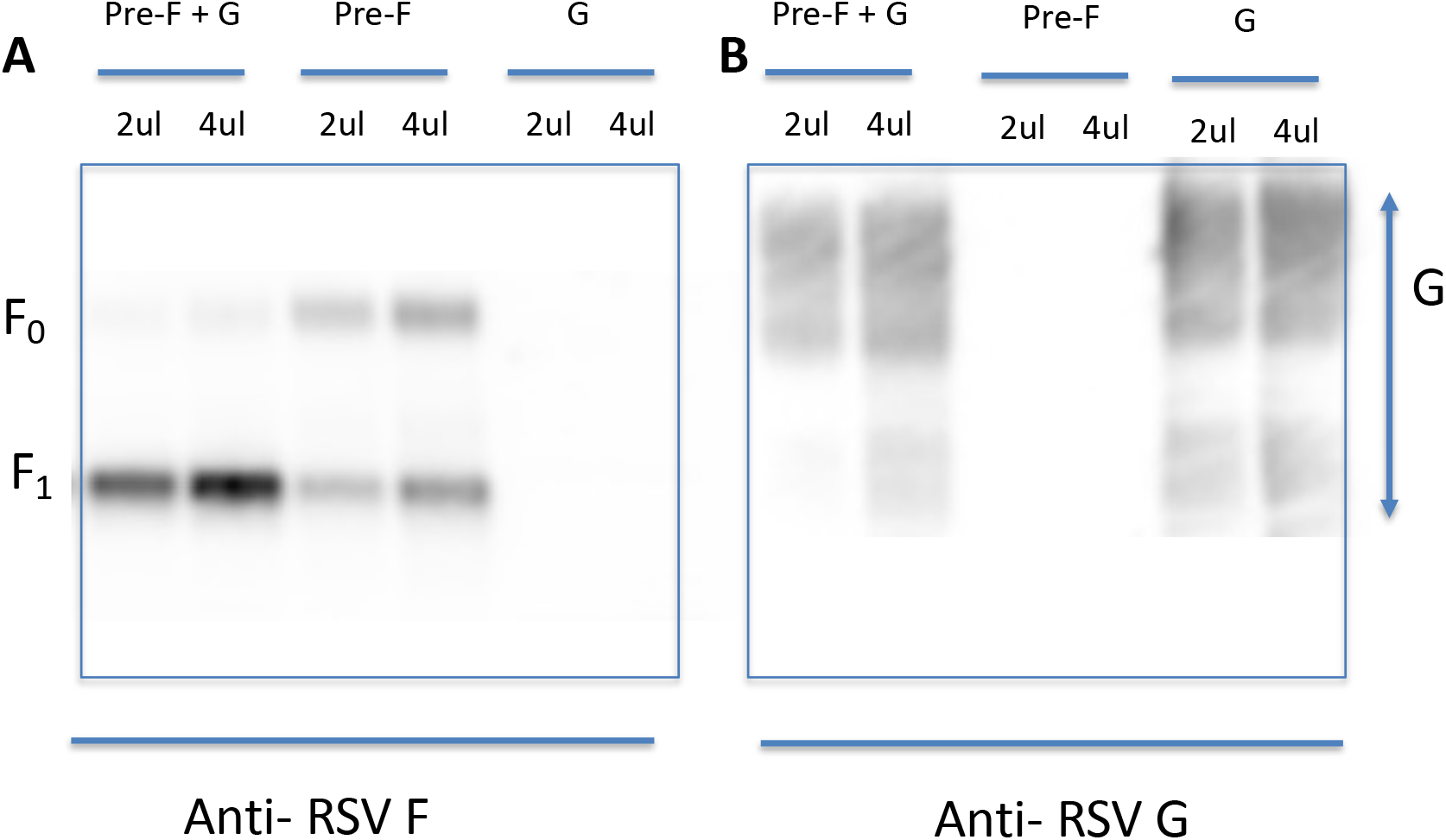
F and G protein content in VLPs. The proteins in the three purified VLPs were detected by Western blot using anti-RSV F (panel A) or anti-RSV G antibodies (panel B). DS Cav1 F proteins in pre-F/F +H/G VLPs, pre-F/F VLPs, and H/G VLPs were detected in 2 and 4 microliters of VLP stocks using anti-RSV-F HR2 polyclonal antibodies. In a separate blot the G proteins in the three VLPs were detected by polyclonal anti-G antibodies.

To assess the antigenic sites on the F and G proteins assembled in the three VLPs, the binding of anti-F protein mAbs to VLPs assembled with F only or with both F and G protein (Figure 2) was compared as was the binding of anti-G protein antibodies to the VLPs assembled with only G protein or both F and G protein (Figure 3). VLPs with equivalent levels of F protein were bound to microtiter wells and then incubated with increasing dilutions of anti-RSV F mAbs representing the major F protein antigenic sites 0, I, II, III, IV and the AM14 binding site (Figure 2). All VLPs bound anti-F mAbs, however, the presence of G protein in the VLPs was associated with differences in binding of anti-F protein mAbs with the exception of binding of mAb specific for site III, MPE8, which bound both VLPs equivalently (Figure 2, panel E). Binding of mAbs to sites I and IV to F/F VLPs and to F/F+H/G VLPs (panels C and F) were different at all dilutions of the mAbs. Binding of mAbs to sites 0 and II (panels A and D), and to theAM14 binding site (Panel B), on both VLPs was similar at most dilutions of antibodies. However, binding to F/F VLPs to these antibodies was lower at saturating antibody concentrations than binding to F/F+H/G VLPs. Thus, in contrast to expectations, the presence of G protein, rather than blocking access of F protein to mAbs, actually enhanced mAb binding to the F protein suggesting that assembly of VLPs with both F and G influenced the conformation of the F protein or accessibility to antibodies.

**Figure 2:**
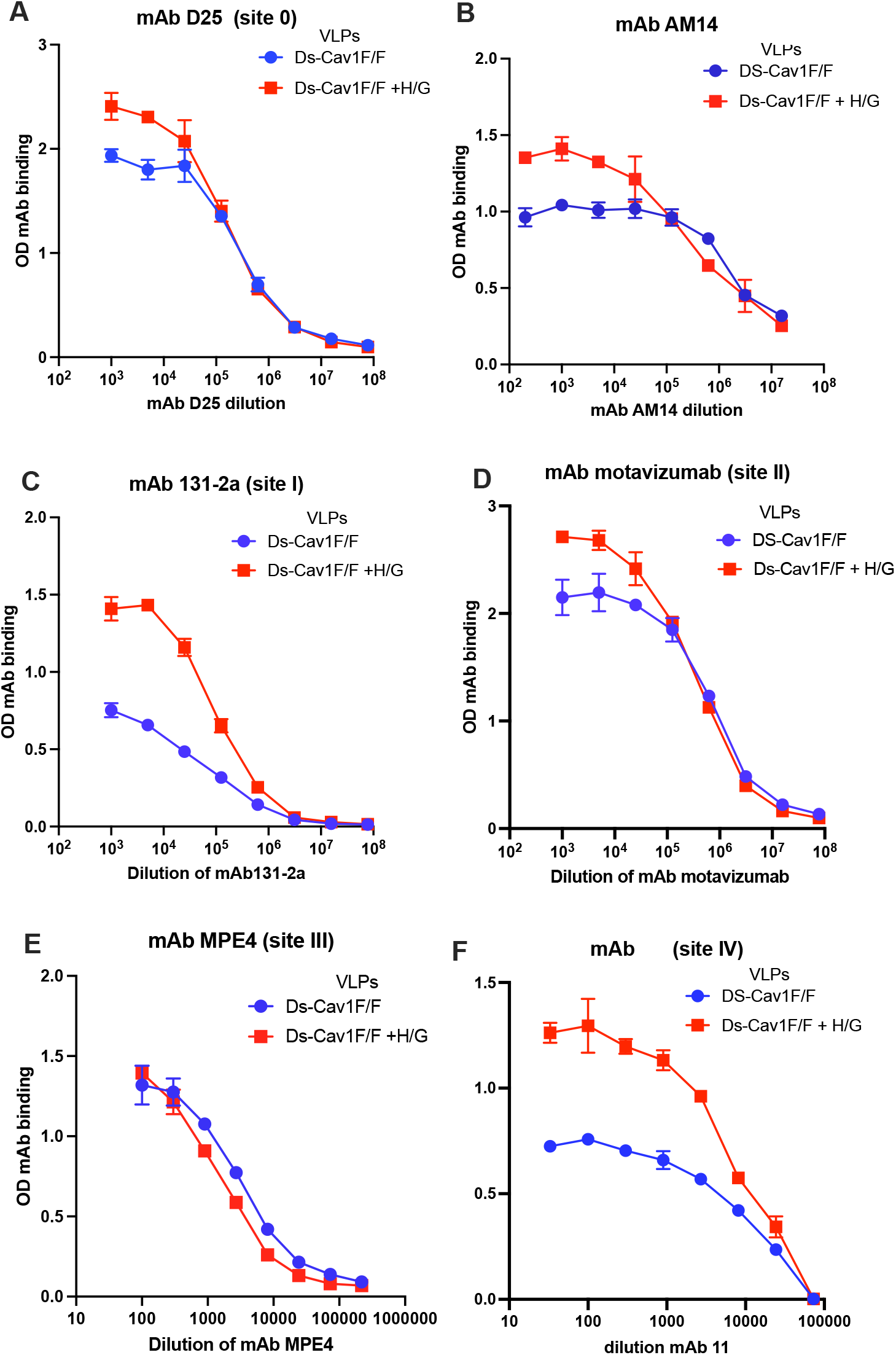
Binding of monoclonal antibodies to purified F/F+ H/G VLPs and F/F VLPs. VLPs containing equivalent levels of RSV F protein were bound to microtiter plates and then incubated with decreasing concentrations of mAb D25 (panel A), AM14 (panel B), mAb 131 −2a (panel C), motavizumab (panel D), mAb MPE8 (panel E), and mAb1243 (panel F). Antibody binding was detected with anti-human IgG or anti-murine IgG coupled with HRP. Results are the average of three separate determinations. The mean and standard deviations of the data from the three determinations at each antibody dilution are shown.

**Figure 3:**
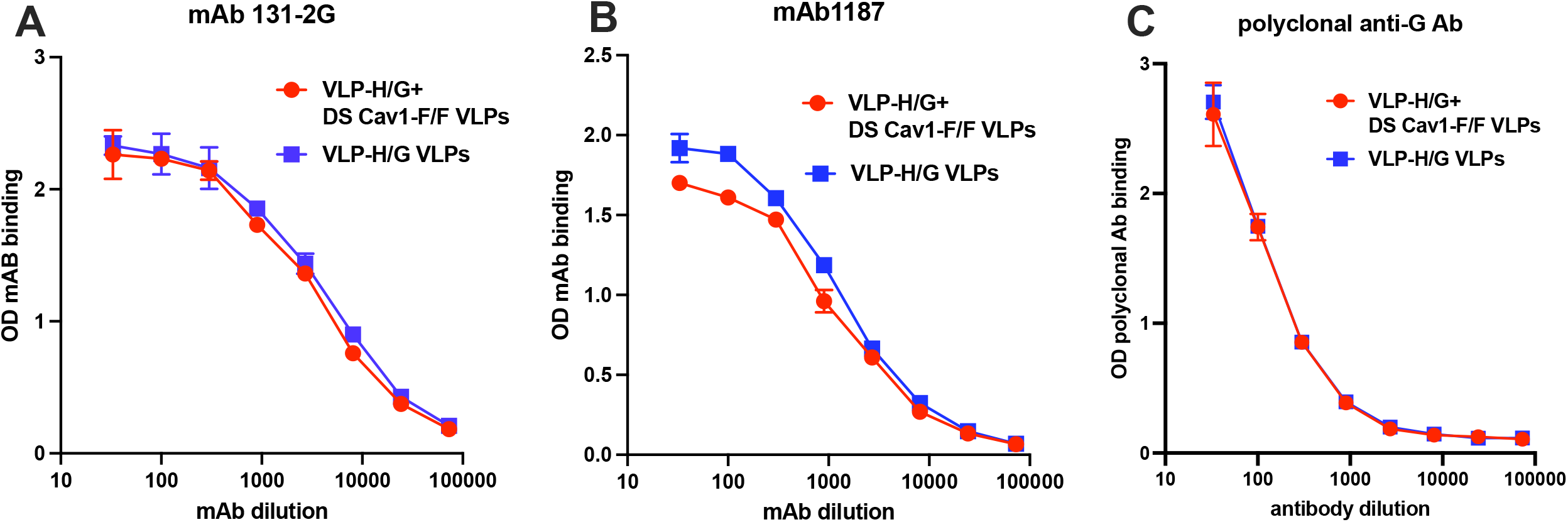
anti-G antibody binding to purified F/F+H/G VLPs and H/G VLPs. VLPs containing equivalent amounts of G protein were bound to microtiter plates and then incubated with decreasing concentrations of mAb 131-2G (panel A), mAb1187 (panel B), or polyclonal anti-G antibody (panel C). Binding of mAbs was detected with anti-human or anti-murine IgG coupled to HRP. Polyclonal Ab binding was detected using anti-rabbit IgG coupled to HRP.

To assess the influence of F protein on antibody binding to G protein, VLPs with equivalent amounts of G protein were bound to microtiter wells and then incubated with two anti-G mAbs or with an anti-G polyclonal antibody. Binding of these antibodies was unaffected by the presence or absence of F protein (Figure 3).

### Immunization protocol

Cotton rats are an accepted animal model for RSV studies of vaccine candidates or inhibitors (33). These animals are much more permissive to RSV replication than mice (33). Furthermore, it has been recently shown that cotton rats express CX3CR1 in their lungs that is very similar to the human equivalent (34). Importantly, it has also been shown that RSV infection of cotton rat lungs is dependent upon CX3CR1 further validating these animals as a model of human infection and for vaccine candidate testing (34).

To compare the immunogenicity of the three VLPs, a group of five 3 to 7-week-old cotton rats were immunized intramuscularly with F/F VLPs containing 5 ug F/F protein. Another group of five animals was immunized with F/F+ H/G VLPs containing 5 ug F/F and 5 ug of H/G. A third group was immunized with H/G VLPs containing 5 ug of H/G protein. A fourth group of 5 cotton rats were mock immunized with PBS. Importantly, all cotton rats were RSV naïve, that is, not previously infected with RSV. Animals were boosted with the same VLPs and the same amount of VLPs on day 28. Sera were collected at days 28, 56, and 60. Animals were challenged with RSV on day 56 and sacrificed on day 60.

### Total anti-pre-F and anti-G antibodies in cotton rat sera

We first measured, by ELISA, the levels of total anti-pre-F and anti-G IgG antibodies induced by the three different VLP immunizations. These assays were accomplished using two different soluble F proteins as ELISA targets. One was the soluble version of the DS-Cav1 mutant F protein. The other was a soluble version of another mutant stabilized, but uncleaved pre-fusion F protein which we have characterized previously, UC-3 pre-fusion F protein (20, 32, 35, 36). Inclusion of the UC-3 F target was because we have detected differences between the two targets in assessing induced anti-RSV antibodies (20, 32, 36).

The presence of G (H/G) in the VLPs did not make a statistically significant difference in the levels of total anti-pre-fusion F IgG antibodies induced using either mutant stabilized pre-F as target (Figure 4, panels A, B). As we have previously noted, however, the levels of total anti-pre- F IgG antibodies detected with the soluble UC-3 F protein target were consistently slightly higher than the levels detected with the soluble DS Cav1 F protein target, but the differences were not statistically significant (36).

**Figure 4:**
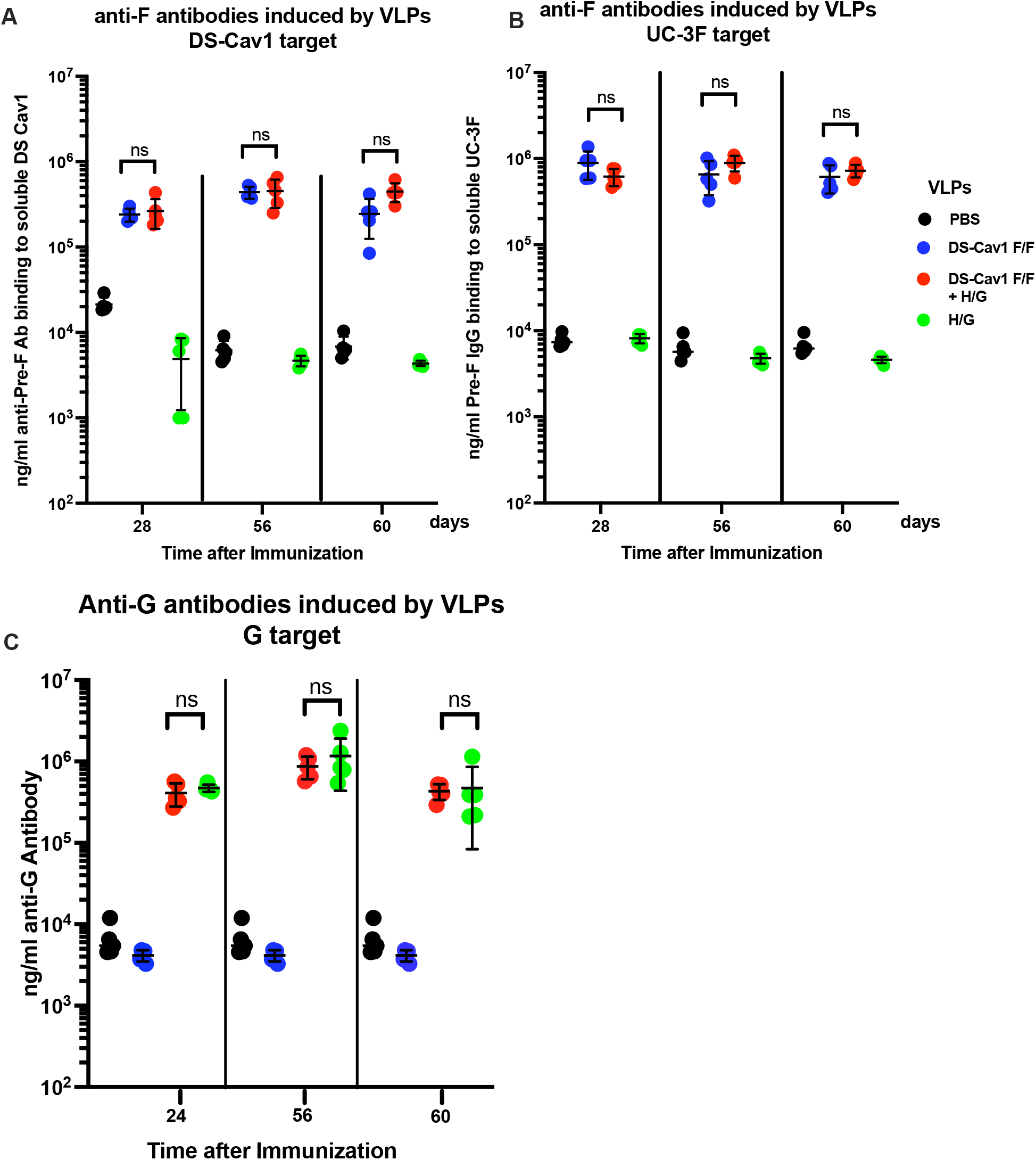
Total anti-pre-F IgG in sera of immunized cotton rats. The concentrations (ng/ml) of anti-pre-F binding IgG in sera of cotton rats immunized with pre- F/F VLPs, pre-F/F+H/G VLPs, or H/G VLPs, as well as mock immunized animals were assessed by ELISA using soluble DS Cav1 F protein as target (panel A), soluble UC-3 F protein as target (panel B), or soluble G protein as target (panel C). Results from sera of individual animals are shown. Shown are titers in each group at 28 days after the prime immunization and times subsequent to the boost at day 28 (days 56 and 60) in individual animals. Black symbols, mock immunization; blue symbols, pre-F/F VLP immunization; red, F/F+H/G VLP immunization; green, H/G VLP immunization. The mean and standard deviations are indicated for each group, shown as black lines. There was no statistically significant difference between in anti-F IgG titers in sera from VLPs containing the F protein. There was no statistically significant difference in anti-G titers in sera from animals immunized with VLPs containing G protein. Sera from H/G VLPs had no detectable anti-F titers above that of mock immunized animals and sera from pre-F/F VLPs had no detectable anti-G titers above that of mock immunized animals.

The levels of anti-G antibodies in sera from animals immunized with VLPs containing only the H/G protein were not significantly different than the levels induced by the F/F+H/G VLPs (Figure 4, panel C).

### Levels of neutralizing antibodies induced by the three VLPs

We next asked if F or G proteins in the VLPs influenced the levels of neutralizing antibodies in the cotton rat sera. The neutralizing antibody (NAb) titers were measured in the classical plaque reduction assay. Because we, and others (37), have previously noted that RSV infections in different cell lines can vary, we accomplished these assays using Vero, Vero E6, A549 p6, and A549 p40 (Figure 5). Surprisingly, sera from animals immunized with F/F VLPs had very low NAb titers titrated in all four cell lines. Sera from animals immunized with H/G VLPs had significantly higher NAb titers but still low compared to titers in sera from animals immunized with F/F+H/G VLPs, which contained high titers of NAb. Thus, assembly of VLPs with both F and G ectodomains significantly impacted the levels of NAb in sera of immunized animals resulting in titers that were between 11 and 12 times higher than the titers in sera from F/F VLP immunization and 3 to 6 times higher than in sera from H/G VLPs immunization. The titers were also higher than the combined titers in sera from F/F VLP and H/G VLP immunizations. This result suggests that the F and G proteins assembled in the same VLP cooperate in some way to induce increased NAb titers.

**Figure 5:**
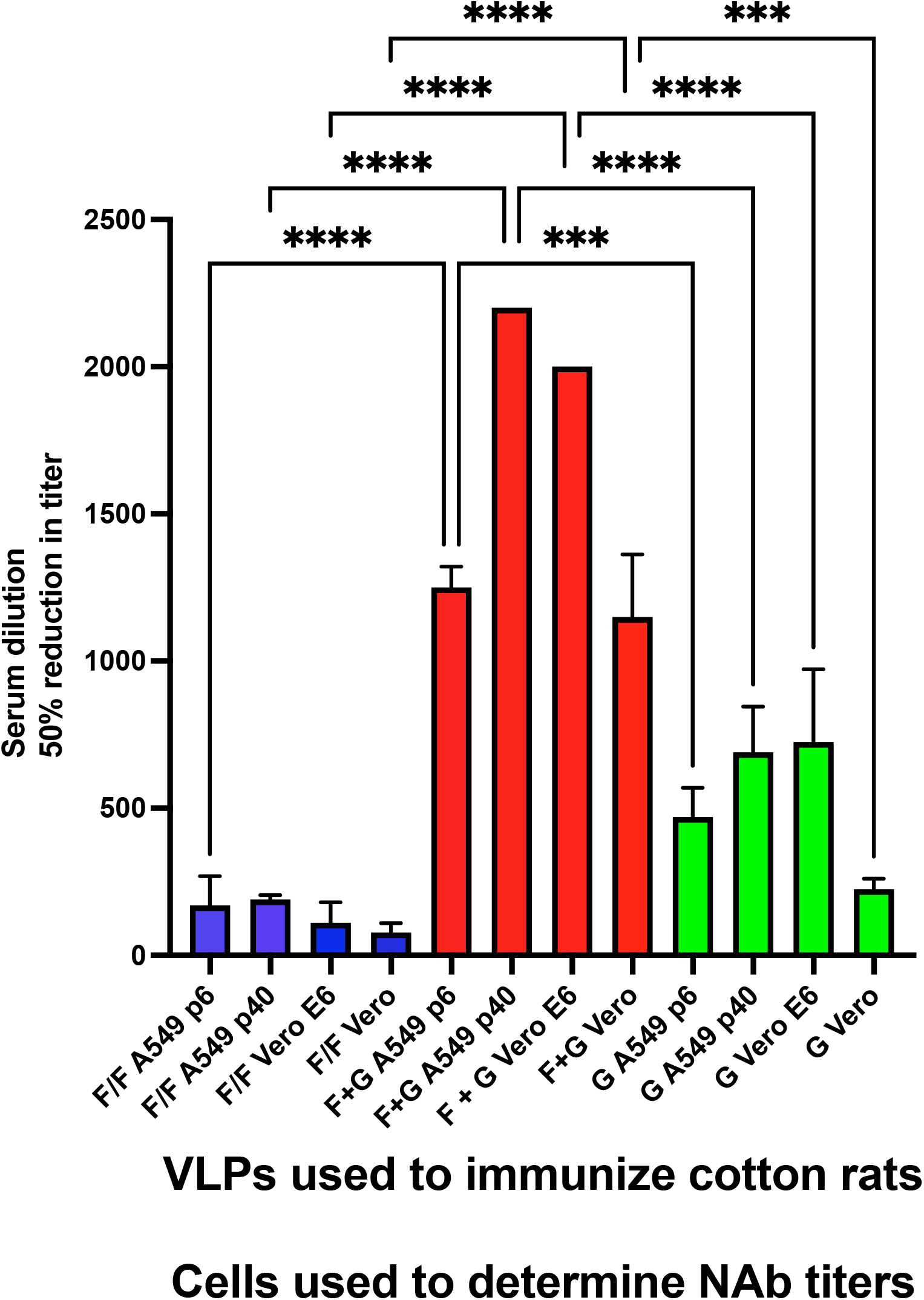
Neutralization titers in cotton rat sera. Sera from each group of animals on day 56 were pooled. NAb titers, determined by the classical plaque reduction assay, were measured using four different cell lines: A549 cells in passage 6 (p6) or passage 40 (p40), Vero cells, or VeroE6 cells. Titers shown are the means and standard deviations of three separate determinations. The statistically significant differences between different groups of animals in titers using the same cells are shown using student t test: p<0.05 *; p<0.005**; p<0.0005***; p<0.00005****.

### Protection of animals from RSV challenge

We next asked if the differences in titers of NAbs antibodies in the sera induced by the three VLPs impacted protection from RSV replication upon challenge of the immunized animals by measuring the levels of virus in the lungs and nasal tissue of the immunized, RSV challenged animals (Figure 6, Panels A and B respectively). As previously reported (19, 21), the lungs of animals immunized with the F/F+H/G VLP immunized animals had no detectable virus and only one animal immunized with H/G VLPs had detectable virus. However, the lungs of the individual animals immunized with F/F VLPs all had detectable virus and the levels were quite variable from animal to animal. Thus, the F/F VLPs did not induce levels protection in the lung comparable to that induced by the other two VLPs.

**Figure 6:**
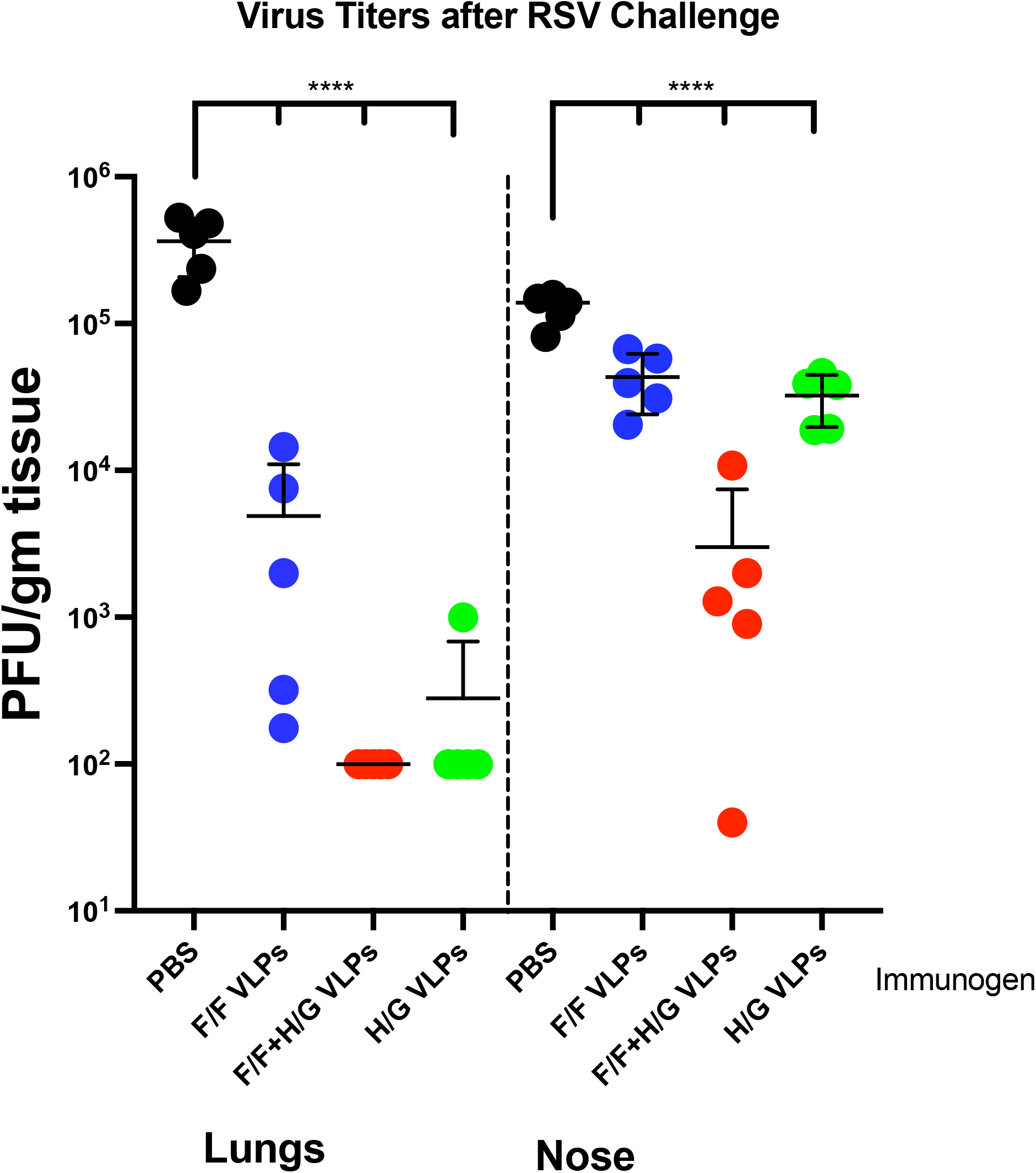
Protection from RSV challenge. Virus titers in lungs and nasal tissue of individual animals after RSV challenge were measured by plaque assay, using HEp2 cells, of different dilutions of homogenized lung or respiratory tissue. Data are shown as plaque forming units (PFU) per gm of tissue. The mean and standard deviation for data from each group are shown (solid lines). Statistical analysis was accomplished using one way ANOVA followed by Dunnett’s multiple comparisons test.

The nose titers of these challenged animals were also measured. As previously noted (21, 22), protection from RSV replication in the upper respiratory tract was weaker than the protection from replication in the lungs after the F/F+H/G VLPs immunization, however protection from upper respiratory infection afforded by these VLPs was better than that afforded by the H/G VLPs or the F/F VLPs.

### Specificities of anti-F antibodies in cotton rat sera

Because the levels of total anti-pre-F binding IgG in sera from cotton rats immunized with the two F/F containing VLPs were the same but the levels of NAb induced by the two VLPs were strikingly different and levels of protection from challenge were different, we assessed the specificities of anti-F antibodies in sera from these animals in order to begin to understand these results. We have previously defined levels of antibodies in polyclonal sera which target specific sites on the F protein by measuring the ng/ml of total anti-pre-F IgG in sera that can block 50 % of the binding of a mAb to a specific F protein epitope, using protocols we have described previously (20, 32). In this way, we can directly compare the levels of antibodies in different sera which target a specific antigenic site on the F protein.

Using this protocol, we measured the levels of antibodies in the three sera specific to epitopes in sites 0, site II, site III, and AM14 binding site, sites all implicated in contributing to induction of high levels of neutralizing antibodies (38). Monoclonal antibodies used were D25 and a humanized 5C4 (both specific to site 0 (27, 39)), MPE8 (site III) (40), AM14 (sometimes identified as site V) (41), and palivizumab (site II) (42, 43). The lower the number of ng/ml of anti-pre-fusion antibodies required to block 50% of the binding of a mAb indicates a higher concentration of antibodies in the sera specific to that site. Using this assay, we found that antibodies specific to F protein sites associated with potent NAb in the sera of animals immunized with F/F+H/G VLPs were much more concentrated than levels in of these antibodies in sera of animals immunized with F/F VLPs (Figure 7). For example, to block 50 % of the binding of 5C4 mAb to F protein required approximately 4000 ng/ml of anti-pre-F IgG induced by F/F+H/G VLPs while approximately 12000 ng of pre-F IgG in sera induced by F/F VLPs were required to block 5C4 mAb binding. The differences in ng/ml required to block D25 binding by the two anti-F containing sera were not as dramatic (15,000ng vs 24,000 ng) but the differences were still statistically significant. Perhaps the differences in concentration of 5C4 and D25 blocking antibodies in the two sera can be attributed to the different binding modes of the two antibodies to site 0 (44). As a control, sera induced by the H/G VLPs did not block any of the mAb binding to F protein (soluble DS-Cav1). These results suggest that the presence of G protein in the VLPs influenced the specificities of anti-F antibodies induced in cotton rats.

**Figure 7:**
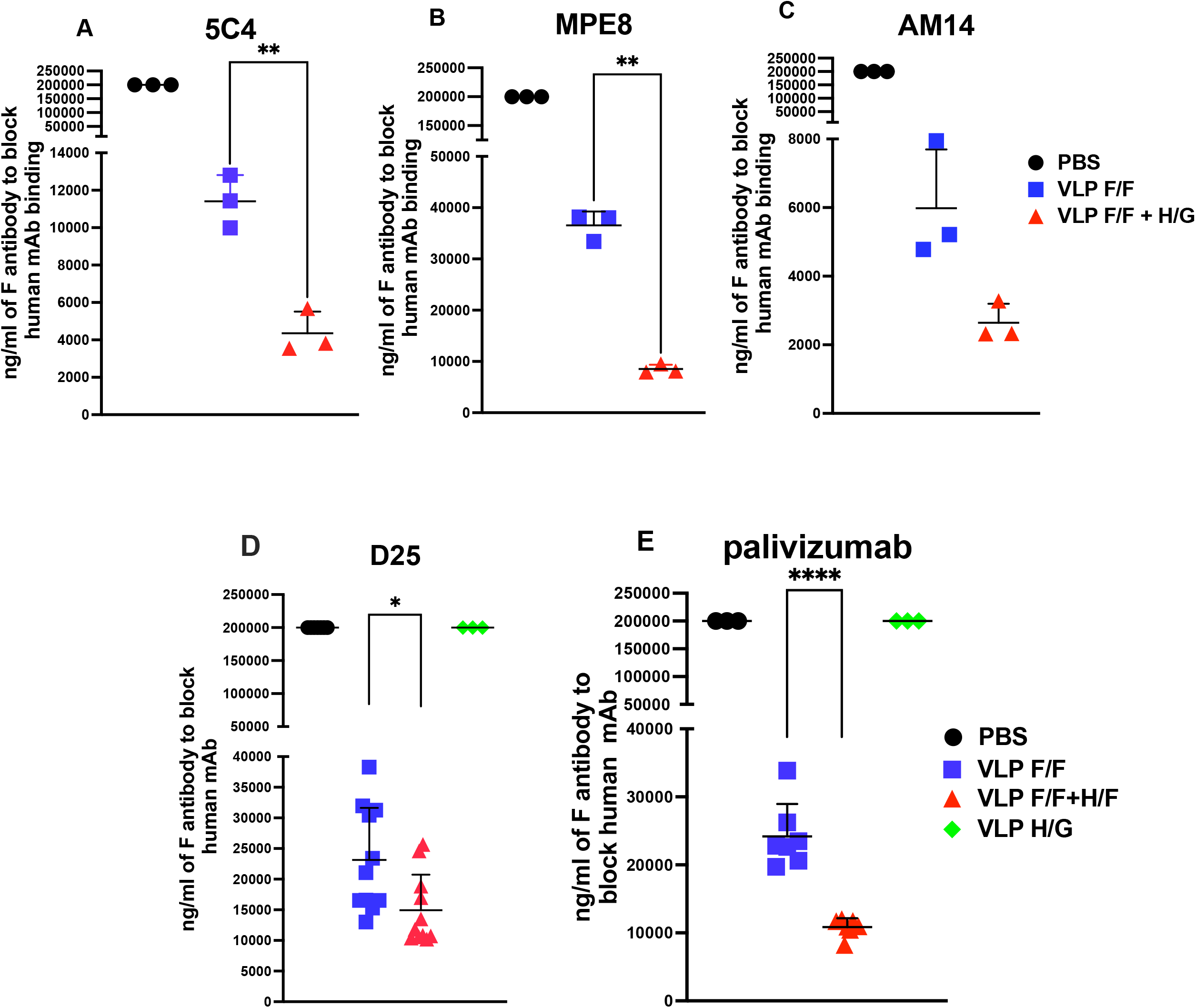
Specificities of antibodies in polyclonal cotton rat sera. Shown are the ng/ml of anti-pre-F IgG in pooled sera from day 56 of each of the four groups of animals that blocked 50% of the binding of different mAb to soluble DS-Cav1 protein. Assay of binding of 5C4 (panel A), MPE8 (panel B), and AM 14 (panel C) were accomplished three times. Assays for D25 mAb (panel D) were accomplished three times in triplicate, assays of palivizumab (panel E) were accomplished two times in triplicate. The results of all assays are shown. Solid lines show the mean and standard deviation of the data. Statistical analysis was accomplished by student t test; p<0.05 *; p<0.005**; p<0.0001****.

## Discussion

This study was designed to address directly the relative importance of F and G proteins in a virus-like particle RSV vaccine candidate. For these studies, we constructed VLPs containing only the mutation-stabilized pre-fusion DS Cav1 F protein (27), VLPs containing only the G protein, and VLPs with both RSV proteins. All three VLPs were assembled with the NP and M core proteins of Newcastle disease virus. Our results showed that assembly of F/F protein with H/G protein enhanced the proteolytic cleavage of the F protein (Figure 1), a result that suggests an influence of G protein on the conformation of the F protein. Furthermore, the VLP binding of most mAbs specific to F protein was enhanced by assembly of F with G protein (Figure 2). While the inefficient cleavage of F protein in F/F VLPs may account for the decreased binding of some anti-F protein mAbs to the F/F VLPs compared to the F/F+H/G VLPs, this explanation seems less likely since we have previously shown that VLPs assembled with an uncleaved, mutation stabilized F protein bound mAbs better than the VLPs containing the cleaved DS Cav1 F protein (32), albeit these results were obtained with VLPs also assembled with the H/G protein. An alternative explanation of the results described here may be due to an impact of the G protein on the conformation of the F protein. We did not detect any influence of the F protein on the conformation of G protein or the anti-G antibodies induced by immunization with G containing VLPs (Figure 3).

Our results also showed that VLPs containing both F and G protein stimulated significantly higher titers of NAb than VLPs containing only F/F or only H/G protein and titers were higher than the sum of NAb titers induced by the F only and G only VLPs (Figure 5). To account for these observations, we quantified the levels of epitope specific antibodies in the polyclonal sera induced by the two F/F containing VLPs by measuring the total ng/ml of anti-pre-F antibodies in the sera that would block the binding of mAb to specific sites on the F protein (Figure 7). In this way, we showed that the presence of G protein in VLPs resulted in increased serum concentrations of antibodies specific for sites associated with inducing high titers of NAb, sites 0, III, and the AM14 binding site, and may account, at least in part, to the higher NAb titers induced by F/F+H/G VLPs compared to that induced by the F/F VLPs. It is also possible that the presence of G protein revealed new epitopes on the F protein.

While results showed significantly different NAb titers induced by F/F VLPs or F+G VLPs, the total anti-pre-F antibodies in the two sera were the same, that is, not affected by the presence of G protein (Figure 4). Anti-pre-F titers were measured using soluble DS Cav1 F protein as target as well as soluble UC-3 pre-F protein. It is well established that many of the epitopes on the two thus far defined conformations of F protein, the pre-fusion F and the post-fusion F, are shared (38, 45). Thus, the anti-pre-F binding IgG in sera likely contains Ab common to epitopes in all different forms of the F protein. The levels of antibodies specific for site 0, site III, and the AM14 binding site may be a minor component of total anti-F serum antibodies and do not affect measures of total pre-F IgG. Alternatively, the F/F+H/G VLPs and F/F VLPs may induce different sets of antibodies with different ratios of antibodies specific for sites associated with induction of potent neutralizing antibody titers.

The very low levels of NAb induced by the F/F VLPs is puzzling given that many RSV vaccine candidates containing only F protein and tested in humans do stimulate NAbs (for example Schmoele-Thoma, et al.(46). How can one, therefore, account for the very low levels of NAbs we observe in cotton rat sera immunized with F/F VLPs? We have previously shown that immunization of RSV naïve cotton rats with soluble DS Cav1 F protein stimulated barely detectable NAbs (Figure 2 in Blanco, et al (22)), a result that is very similar to our findings here with F/F VLPs. However, in this earlier report, we also showed that immunization of animals previously infected with RSV, RSV primed, followed with immunization with soluble DS Cav1 F protein did stimulate NAbs (Figure 2 in Blanco, et al (22)) albeit still lower than titers induced by the F/F+H/G VLPs. We proposed that the different results obtained in RSV naïve animals and RSV primed animals were due to the establishment of memory to RSV epitopes by the prior RSV infection and that the soluble F protein could activate memory responses established in the previous RSV infections, but that the soluble F protein in naïve animals was less effective in stimulating de novo responses thus resulting in little detectable NAbs. The vast majority of humans have been infected with RSV by 2 to 5 years of age and many have experienced multiple infections (24, 25). Thus, we suggest that NAbs stimulated by soluble F protein vaccine candidates in humans are due to stimulation of memory responses established by previous RSV infections.

In contrast to soluble F protein, we have shown here and previously that F/F+H/G VLPs can stimulate neutralizing antibodies in RSV naïve animals (22). We also previously showed that immunization of RSV primed animals with F/F+H/G VLPs stimulated significantly higher NAbs than immunization of RSV naïve animals suggesting that these VLPs can also activate memory responses (22). However, a second RSV infection of RSV primed animals only resulted in a very weak stimulation of anti-RSV antibodies (22, 23, 32), which may account for the susceptibility of humans to multiple RSV infections throughout their lifespan.

Indirect evidence presented here of an influence of the G protein on the properties of the F protein and on induction of NAbs would most logically be explained by an interaction of the two proteins in VLPs or during the folding of the proteins prior to assembly into VLPs. Indeed, there are two reports of interactions between the F and G proteins of the Pneumoviridae RSV (47) and metapneumovirus (48). This issue has been extensively studied for many of the viruses in the closely related Paramyxoviradae family (for example (49-56)). Currently the most widely accepted view concerning paramyxovirus glycoproteins is that interactions are weak and/or transient and occur either before or after the binding of the attachment protein to its receptor, depending upon the virus (53). However, complexes between the fusion and attachment paramyxovirus proteins have been difficult to conclusively demonstrate and their existence has recently been challenged by Wong et al. (57). The F and G interactions in Pneumoviridae have not been studied in detail thus the mechanisms involved in the influence of G protein on F protein antibody responses and F protein interactions with mAbs in the presence of G protein reported here remain to be elucidated.

In summary, our results here suggest that inclusion of both F and G protein in vaccine candidates that allow both proteins to directly interact will result in higher titers of NAbs and likely improved protection from RSV infections. Formulation of vaccine candidates for different populations, RSV experienced adults vs RSV naïve infants and young children, should consider the importance of G protein in those candidates relative to efficacy as well as safety.

## Methods

### Preparation, characterization, and validation of VLP stocks

VLPs used as immunogens were based on the core proteins of Newcastle disease virus (NDV) M and NP proteins and contained the RSV F and G glycoproteins (19, 29, 31). The RSV proteins were assembled into the VLPs as chimera proteins with the sequences of the ectodomain of RSV F and G glycoproteins (sequences from RSV strain A2) fused to the transmembrane and cytoplasmic domains of the NDV F and HN proteins, respectively to create the F/F or the H/G chimera proteins. The F/F protein contained the pre-fusion stabilizing DS Cav1 mutations (27) and the foldon sequence (28) was inserted between the F protein ectodomain and the NDV F protein transmembrane domain to further stabilize the pre-fusion form of the protein.

Three different VLPs were prepared. One VLP contained only the F/F chimera protein. A second VLP was assembled with only the H/G chimera protein. A third VLP was assembled with both the DS Cav1 F/F and the H/G chimera protein. The VLPs were prepared by transfecting avian cells (ELL-0, ATCC CRL-12203) with cDNAs encoding the NDV M and NP proteins as well as the G protein chimera or the F chimera protein or both the F and G chimera proteins. VLPs released into the cell supernatant were purified as previously described (30) and the F and G protein content of purified VLPs were quantified by Western blots and by monoclonal antibody (mAb) binding to the VLPs as previously described (22, 32). VLP stocks were adjusted for equivalent levels of F protein and G protein (22, 32). The pre-fusion conformation of the F protein in the VLPs was validated by assessing the binding of mAbs specific to the pre-fusion form of the F protein to the VLPs as previously reported (22, 32).

### Preparation of soluble DS Cav1 F protein, UC3 soluble F protein, or the soluble G protein

Expi293F cells were transfected with cDNAs encoding the soluble DS-Cav1 pre-F protein, the soluble UC-3 F protein, or the soluble G protein. At six days post transfection, total cell supernatants were collected, cell debris removed by centrifugation, and the soluble polypeptides were purified on columns using the His tag and then the streptavidin tag (22, 58). Purified soluble DS-Cav1 pre-F protein or UC-3 F protein and soluble G protein efficiently bound to pre-fusion specific mAbs AM14 and D25 or mAb specific to the RSV G protein (32).

### Quantification of NP, M, H/G and VLP associated F proteins or soluble F proteins

For Western blots, proteins were resolved on 8% Bis-Tris gels (NuPage, ThermoFisher/Invitrogen). Quantifications of NP, M, H/G proteins, and F/F proteins in VLPs or in soluble F protein or G preparations were accomplished after their separation in polyacrylamide gels followed by silver staining (Pierce Silver Stain, ThermoFisher) or Western blots of the proteins in parallel with protein standards as previously described (22, 58).

### Monoclonal antibody (mAb) binding to purified VLPs

VLPs containing equivalent amounts of F protein or equivalent amounts of G protein (determined by Western blot) were added to microtiter wells and incubated overnight at 4^0^C. Increasing dilutions of different mAbs were added to the wells, incubated for 2 hours, removed, and the wells washed with PBS. Goat anti-human IgG or anti-murine IgG coupled to HRP was added to each well and incubated for 2 hours at room temperature, removed, and the plate washed extensively with PBS. Bound HRP was detected using TMB (3,3’.5.5’- tetramethylbenzidin, ThermoFisher 34028, Walthan MA, USA) and the reaction was stopped with 2N sulfuric acid. Color was read in the SpectraMax Pro Plate Reader (Molecular Devices) using SoftMax Pro software. Results are expressed as optical density (OD).

### ELISA

For determination of anti-pre-F protein IgG antibody titers in sera, wells of microtiter plates (ThermoFisher/Costar) were coated with purified soluble DS-Cav1 F protein or soluble UC-3 F protein (30 ng/well) and incubated overnight at 4°C, then blocked with 2% BSA for 16 hours at 4^0^C. Different dilutions of sera, in PBS-2% BSA and 0.05% Tween, were added to each well and incubated for 2 hours at room temperature. Wells were then washed with PBS, incubated with chicken anti-cotton rat IgG antibody (Abnova PAB29753) coupled to HRP, and incubated for 1.5 hours at room temperature. Wells were washed and bound HRP was detected using TMB as described above. Amounts of IgG (ng/ml) in each dilution were calculated using a standard curve generated using defined amounts of purified cotton rat IgG, as previously described (22).

### RSV Neutralization

RSV was grown in HEp-2 (ATCC.CCL-23) cells, and RSV plaque assays were accomplished on Vero, Vero E6, A549, passage 6, or A549, passage 40, as previously described (22, 58). Antibody neutralization assays in a plaque reduction assay have been previously described (58). Neutralization titer was defined as the reciprocal of the dilution of serum that reduced virus titer by 50%.

### Animals

*Sigmodon hispidus* cotton rats (CR) were obtained from Envigo (Indianapolis, IN, USA). All studies were conducted under applicable laws and guidelines and after approval from Merck, Inc. Institutional Animal Care and Use Committee. Animals were housed in large polycarbonate cages and fed a standard diet of rodent chow and water ad libitum. Animals were pre-bled before inclusion in the study to rule out the possibility of pre-existing antibodies against RSV.

### Experimental design

Groups of five animals, 3-7 weeks of age, were immunized intramuscularly with F/F VLPs containing 5 ug F/F protein. Another group of five animals was immunized with F/F+ H/G VLPs containing 5 ug F/F and 5 ug of H/G. A third group was immunized with H/G VLPs containing 5 ug of H/G protein. A fourth group of 6 cotton rats were mock immunized with PBS. Importantly, all cotton rats were RSV naïve, that is, not previously infected with RSV. Animals were boosted with the same VLPs and the same amount of VLPs on day 28. Sera were collected at days 28, and 56. Animals were challenged with RSV by intranasal inoculation on day 56 and sacrificed on day 60. Lungs and nasal tissue were harvested for titration of virus. Virus titers in lungs and nasal tissue of individual animals after RSV challenge were measured by plaque assay, using HEp2 cells, of different dilutions of homogenized lung or nasal tissue.

### Cells, virus, and antibodies

The cell line used to produce VLPs was ELL0 (ATCC CRL-12203). Cell lines used for plaque assays were A549, Vero, and VeroE6, all obtained from ATCC. Two passages of A549 were used: passage 6 after recovery from frozen storage and passage 40 after recovery from storage. The virus strain used for RSV challenge experiments was A2 and the cDNA clones of RSV sequences were derived from the RSV A2 strain. Antibodies used for Western blots were anti-RSV HR2 raised in rabbits using as antigen the RSV F HR2 domain, as previously described (29). Antibody used in plaque assays was mAb131-2a (MAB8599, Millipore). A polyclonal anti-G antibody was obtained from ThermoFisher. Monoclonal antibodies specific to F protein D25, palivizumab, AM14, motavizumab, 131-2a, mAb 1243 and G protein specific mAb 131-2G, mAb1187 were obtained from B. Graham, J. Blanco, J. McLellan, and J. Beeler. Monoclonal antibodies MPE8 and 5C4 were humanized versions of anti-F protein specific murine antibodies. Secondary antibodies against goat, mouse, human, and rabbit IgG were purchased from Sigma.

### Statistical analysis

Statistical analyses were accomplished using ANOVA (followed by post-hoc Tukey HSD or Sidák multiple comparison test) or unpaired student-*t* test of data (indicated in the figure legends) all of which were accomplished using Graph Pad Prism 9 software.

## Acknowledgements

This project was supported by National Institutes of Health grant AI043896 awarded to T Morrison and by Merck & Co, Inc. We thank Drs J. Beeler, J. McLellan, B. Graham, and J. Blanco for generous gifts of antibodies.

## Contributions

L. M. C.: Prepared the VLPs, validated the VLPs, Western blots, performed ELISA on VLPs, measured IgG in CR sera, neutralization assays of CR sera, measurements of blocking of mAb binding to targets by serum

L.B.: Performed the in-vivo study

Z.W: Performed viral titers in lung and nose

T.G. M., L.Z., E.D.: Experiment conception / manuscript review

T.G. M.: wrote the manuscript

## Literature Cited

1. Shi T, McAllister DA, O’Brien KL, Simoes EAF, al. e. 2017. Global, regional, and national disease burden estimates of acute lower respiratory infections due to respiratory syncytial virus in young children in 2015: a systematic review and modelling study. The Lancet 390:946–958.

2. Nair H, Nokes DJ, Gessner BD, Dherani M, Madhi SA, Singleton RJ, O’Brien KL, Roca A, Wright PF, Bruce N, Chandran A, Theodoratou E, Sutanto A, Sedyaningsih ER, Ngama M, Munywoki PK, Kartasasmita C, Simoes EAF, Rudan I, Weber MW, Campbell H. 2010. Global burden of acute lower respiratory infections due to respiratory syncytial virus in young children: a systematic review and meta-analysis. The Lancet 375:1545–1555.

3. Thompson W, Shay DK, Weintraub E, et al. 2003. Mortality associated with influenza and respiratory syncytial virus in the United States. JAMA 289:179–186.

4. Raboni SM, Nogueira MB, Tsuchiya LR, Takahashi GA, Pereira LA, Pasquini R. 2003. Respiratory tract viral infections in bone marrow transplant patients. Transplant 76:142–146.

5. Han LL, Alexander JP, Anderson LJ. 1999. Respiratory syncytial virus pneumonia among the elderly: an assessment of disease burden. J Infect Dis 179:25–30.

6. Falsey AR, Walsh EE. 2000. Respiratory syncytial virus infection in adults. Clin Microbiol Rev 13:371–384.

7. Falsey AR, Hennessey PA, Formica MA, Cox C, Walsh EE. 2005. Respiratory syncytial virus infection in elderly and high-risk adults. N Engl J Med 352:1749–1759.

8. Shan J, Britton PN, King CL, Booy R. 2021. The immunogenicity and safety of respiratory syncytial virus vaccines in development: A systematic review. Influenza and Other Respiratory Viruses 15:539–551.

9. Melero JA, Moore ML. 2013. Influence of Respiratory Syncytial Virus Strain Differences on Pathogenesis and Immunity, p 59–82. In Anderson LJ, Graham BS (ed), Challenges and Opportunities for Respiratory Syncytial Virus Vaccines, vol 372. Springer-Verlag, Berlin Heidelberg.

10. Anderson LJ, Jadhao SJ, Paden CR, Tong S. 2021. Functional Features of the Respiratory Syncytial Virus G Protein. Viruses 13:1214.

11. Johnson SM, McNally BA, Ioannidis I, Flano E, Teng MN, Oomens AG, Walsh EE, Peeples ME. 2015. Respiratory Syncytial Virus Uses CX3CR1 as a Receptor on Primary Human Airway Epithelial Cultures. PLoS Pathog 11:e1005318.

12. Choi Y, Mason CS, Jones LP, Crabtree J, Jorquera PA, Tripp RA. 2012. Antibodies to the Central Conserved Region of Respiratory Syncytial Virus (RSV) G Protein Block RSV G Protein CX3C-CX3CR1 Binding and Cross-Neutralize RSV A and B Strains. Viral Immunol 25:193–203.

13. Chirkova T, Lin S, Oomens AGP, Gaston KA, Boyoglu-Barnum S, Meng J, Stobart CC, Cotton CU, Hartert TV, Moore ML, Ziady AG, Anderson LJ. 2015. CX3CR1 is an important surface molecule for respiratory syncytial virus infection in human airway epithelial cells. J of Gen Virol 96:2543–2556.

14. Tripp RA, Jones LP, Haynes LM, Zheng H, Murphy PM, Anderson LJ. 2001. CX3C chemokine mimicry by respiratory syncytial virus G glycoprotein. Nat Immunol 2:732–738.

15. Boyoglu-Barnum S, Todd SO, Chirkova T, Barnum TR, Gaston KA, Haynes LM, Tripp RA, Moore ML, Anderson LJ. 2015. An anti-G protein monoclonal antibody treats RSV disease more effectively than an anti-F monoclonal antibody in BALB/c mice. Virology 483:117–125.

16. Anderson LJ, P. B, Hierholzer JC. 1988. Neutralization of respiratory syncytial virus by individual and mixtures of F and G protein monoclonal antibodies. J Virol 62:4232–4238.

17. Hallak LK, Spillmann D, Peeples ME. 2000. Glycosaminoglycan sulfation requirements for respiratory syncytial virus infection. Journal of Virology 74:10508–10513.

18. Hallak LK, Collins PL, Knudson W, Peeples ME. 2000. Iduronic acid-containing glucosaminoglycans on target cells are required for efficient respiratory syncytial virus infection. Virology 271:264–275.

19. Murawski MR, McGinnes LW, Finberg RW, Kurt-Jones EA, Massare M, Smith G. 2010. Newcastle disease virus-like particles containing respiratory syncytial virus G protein induced protection in BALB/c mice with no evidence of immunopathology. J Virol 84:1110–1123.

20. Cullen LM, Boukhvalova MS, Blanco JCG, Morrison TG. 2020. Comparisons of Antibody Populations in Different Pre-Fusion F VLP-Immunized Cotton Rat Dams and Their Offspring. Vaccines 8:133–148.

21. Cullen LM, Blanco JCG, Morrison TG. 2015. Cotton rat immune responses to virus-like particles containing the pre-fusion form of respiratory syncytial virus fusion protein. J Transl Med 13:1–13.

22. Blanco JCG, Pletneva LM, Cullen L, Otoa RO, Patel MC, Fernando LR, Boukhvalova MS, Morrison TG. 2018. Efficacy of a respiratory syncytial virus vaccine candidate in a maternal immunization model. Nature Comm 9:1904–1914.

23. Cullen LM, Schmidt MR, Morrison TG. 2019. Effect of Previous Respiratory Syncytial Virus Infection on Murine Immune Responses to F and G Protein-Containing Virus-Like Particles. J Virol 93:e00087–00019.

24. Hall CB, Simoes EAF, Anderson LJ. 2013. Clinical and Epidemiologic Features of Respiratory Syncytial Virus, p 39–58. In Anderson LJ, Graham BS (ed), Challenges and Opportunities for Respiratory Syncytial Virus Vaccines, vol 372. Springer, Heidelberg, New York, Dordrecht, Londaon.

25. Hall CB, Walsh EE, Long CE, Schnabel KD. 1991. Immunity to and frequency of reinfections with respirtory syncytial virus. J Infect Dis 163:693–698.

26. Bachmann MF, Jennings GT. 2010. Vaccine delivery: a matter of size, geometry, kinetics and molecular patterns. Nat Rev Immunol 10:787–796.

27. McLellan JS, Chen M, Joyce MG, Sastry M, Stewart-Jones Gbe, Yang Y. 2013. Structure-based design of a fusion glycoprotein vaccine for respiratory syncytial virus. Science 342:592–598.

28. Frank S, Kammerer RA, Mechling D, Schulthess T, Landwehr R, Bann J. 2001. Stabilization of short collagen-like triple helices by protein engineering. J Mol Biol 308:1081–1089.

29. McGinnes LW, Gravel KA, Finberg RW, Kurt-Jones EA, Massare MJ, Smith G. 2011. Assembly and immunological properties of Newcastle disease virus-like particles containing the respiratory syncytial virus F and G proteins. J Virol 85:366–377.

30. McGinnes LW, Morrison TG. 2013. Newcastle Disease Virus-Like Particles: Preparation, Purification, Quantification, and Incorporation of Foreign Glycoproteins, Current Protocols in Microbiology doi:10.1002/9780471729259.mc1802s30. John Wiley & Sons, Inc.

31. McGinnes L, Schmidt MR, Kenward SA, Woodland RT, Morrison TG. 2015. Murine Immune Responses to Virus-Like Particle-Associated Pre- and Postfusion Forms of the Respiratory Syncytial Virus F Protein. J Virol 89:6835–6847.

32. Cullen ML, Schmidt RM, Torres MG, Capoferri AA, Morrison GT. 2019. Comparison of Immune Responses to Different Versions of VLP Associated Stabilized RSV Pre-Fusion F Protein. Vaccines 7:21–41.

33. Boukhvalova MS, Blanco J. 2013. The cotton rat Sigmondon Hispidus model of respiratory syncytial virus infection. In Anderson LJ, Graham BS (ed), Challenges and Opportunities for Respiratory Syncytial Virus Vaccines. Springer-Verlag, Berlin Heidelberg.

34. Green G, Johnson SM, Costello H, Brakel K, Harder O, Oomens AG, Peeples ME, Moulton HM, Niewiesk S, Heise MT. 2021. CX3CR1 Is a Receptor for Human Respiratory Syncytial Virus in Cotton Rats. J Virol 95:e00010–00021.

35. Blanco JCG, Fernando LR, Zhang W, Kamali A, Boukhvalova MS, McGinnes-Cullen L, Morrison TG. 2019. Alternative Virus-Like Particle-Associated Prefusion F Proteins as Maternal Vaccines for Respiratory Syncytial Virus. J Virol 93:e00914–00919.

36. Blanco J, Cullen L, Kamali A, Sylla F, Boukhvalova MS, Morrison TG. 2021. Evolution of protection after maternal immunization for respiratory syncytial virus in cotton rats. PLOS Pathog 17:e1009856.

37. Rajan A, Piedra F-A, Aideyan L, McBride T, Robertson M, Johnson HL, Aloisio GM, Henke D, Coarfa C, Stossi F, Menon VK, Doddapaneni H, Muzny DM, Cregeen SJJ, Hoffman KL, Petrosino J, Gibbs RA, Avadhanula V, Piedra PA, Heise MT. 2022. Multiple Respiratory Syncytial Virus (RSV) Strains Infecting HEp-2 and A549 Cells Reveal Cell Line-Dependent Differences in Resistance to RSV Infection. J Virol 96:e01904–01921.

38. Ngwuta JO, Chen M, Modjarrad K, Joyce MG, Kanekiyo M, Kumar A, Yassine HM, Moin SM, Killikelly AM, Chuang G-Y, Druz A, Georgiev IS, Rundlet EJ, Sastry M, Stewart-Jones Gbe, Yang Y, Zhang B, Nason MC, Capella C, Peeples ME, Ledgerwood JE, McLellan JS, Kwong PD, Graham BS. 2015. Prefusion F–specific antibodies determine the magnitude of RSV neutralizing activity in human sera. Science Transl Med 7:309ra162.

39. McLellan JS, Chen M, Leung S, Graepel KW, D. X, Yang Y, Zhou T, Baxa U, Yasuda E, Beaumont T, Kumar A, Modjarrad K, Zheng Z, Zhao M, Xia N, Kwong PD, Graham BS. 2013. Structure of RSV Fusion Glycoprotein Trimer Bound to a Prefusion-Specific Neutralizing Antibody. Science 340:1113–1117.

40. Corti D, Bianchi S, Vanzetta F, Minola A, Perez L, Agatic G, Guarino B, Silacci C, Marcandalli J, Marsland BJ, Piralla A, Percivalle E, Sallusto F, Baldanti F, Lanzavecchia A. 2013. Cross-neutralization of four paramyxoviruses by a human monoclonal antibody. Nature 501:439–443.

41. Gilman MSA, Moin SM, Mas V, Chen M, Patel NK, Kramer K, Zhu Q, Kabeche SC, Kumar A, Palomo C, Beaumont T, Baxa U, Ulbrandt ND, Melero JA, Graham BS, McLellan JS. 2015. Characterization of a Prefusion-Specific Antibody That Recognizes a Quaternary, Cleavage-Dependent Epitope on the RSV Fusion Glycoprotein. PLoS Pathog 11:e1005035.

42. Parnes C, Guillermin R, Habersang R, Nicholas P, Chawla V, Kelly T, Fishbein J, others. 2003. Palivizumab prophylaxis of respiratory syncytial virus disease in 2000-2001: results from the Palivizumab outcomes registry. Pediatr Pulmonol 35:484–489.

43. Mousa JJ, Sauer MF, Sevy AM, Finn JA, Bates JT, Alvarado G, King HG, Loerinc LB, Fong RH, Doranz BJ, Correia BE, Kalyuzhniy O, Wen X, Jardetzky TS, Schief WR, Ohi MD, Meiler J, Crowe JE. 2016. Structural basis for nonneutralizing antibody competition at antigenic site II of the respiratory syncytial virus fusion protein. Proc Natl Acad of Sci USA 113:E6849–E6858.

44. Tian D, Battles MB, Moin SM, Chen M, Modjarrad K, Kumar A, Kanekiyo M, Graepel KW, Taher NM, Hotard AL, Moore ML, Zhao M, Zheng Z-Z, Xia N-S, McLellan JS, Graham BS. 2017. Structural basis of respiratory syncytial virus subtype-dependent neutralization by an antibody targeting the fusion glycoprotein. Nature Comm 8:1877.

45. Smith G, Raghunandan R, Wu Y, Liu Y, Massare M, Nathan M, Zhou B, Lu H, Boddapati S, Li J, Flyer D, Glenn G. 2012. Respiratory Syncytial Virus Fusion Glycoprotein Expressed in Insect Cells Form Protein Nanoparticles That Induce Protective Immunity in Cotton Rats. PLoS ONE 7:e50852.

46. Schmoele-Thoma B, Zareba AM, Jiang Q, Maddur MS, Danaf R, Mann A, Eze K, Fok-Seang J, Kabir G, Catchpole A, Scott DA, Gurtman AC, Jansen KU, Gruber WC, Dormitzer PR, Swanson KA. 2022. Vaccine Efficacy in Adults in a Respiratory Syncytial Virus Challenge Study. N Engl J of Med 386:2377–2386.

47. Low K-W, Tan T, Ng K, Tan B-H, Sugrue R. 2008. The RSV F and G glycoproteins interact to form a complex on the surface of infected cells. BBRC 366:308–313.

48. Loo L, Jumat M, Fu Y, Ayi T, Wong P, Tee N, Tan B-H, Sugrue R. 2013. Evidence for the interaction of the human metapneumovirus G and F proteins during virus-like particle formation.. Virol J 10:294–307.

49. Bose S JT, Lamb RA. 2015. Timing is everything: Fine-tuned molecular machines orchestrate paramyxovirus entry. Virology 479:518–531.

50. Plemper RK, Hammond AL, Gerlier D, Fielding AK, Cattaneo R. 2002. Strength of envelope protein interaction modulates cytopathicity of measles virus. J Virol 76:5051–5061.

51. Bossart K, Wang L, Flora M, Kb C, Lam S, Eaton B, Broder C. 2002. Membrane Fusion Tropism and Heterotypic Functional Activities of the Nipah Virus and Hendra Virus Envelope Glycoproteins. J Virol 76:11186–11198.

52. Porotto M, Murrell M, Greengard O, Moscona A. 2003. Triggering of human parainfluenza virus 3 fusion protein (F) by the hemagglutinin-neuraminidase (HN) protein: an HN mutation diminishes the rate of F activation and fusion. J Virol 77:3647–3654.

53. Bossart KN, Fusco DL, Cc. B. 213. Paramyxovirus entry. Adv Exp Med Biol 790:95–127.

54. Yao Q, Hu X, Compans R. 1997. Association of the parainfluenza virus fusion and hemagglutinin-neuraminidase glycoproteins on cell surfaces. J Virol 71:650–656.

55. Hu X, Ray R, Compans RW. 1992. Functional interactions between the fusion protein and hemagglutinin-neuraminidase of human parainfluenza viruses. J Virol 66:1528–1534.

56. Nadine A, Melinda B, Mislay A, Claes Ö, Branka H, Georg H, Jürgen S-S, Marc V, Andreas Z, Richard KP, Philippe P. 2013. Mechanism for Active Membrane Fusion Triggering by Morbillivirus Attachment Protein. J Virol 87:314–326.

57. Wong JJ, Chen Z, Chung JK, Groves JT, Jardetzky TS. 2021. EphrinB2 clustering by Nipah virus G is required to activate and trap F intermediates at supported lipid bilayer–cell interfaces. Science Adv 7:eabe1235.

58. Cullen LM, Schmidt MR, Morrison TG. 2017. The importance of RSV F protein conformation in VLPs in stimulation of neutralizing antibody titers in mice previously infected with RSV. Human Vaccines & Immunotherapeutics doi:10.1080/21645515.2017.1329069:1–10.

